# How inhibitory neurons increase information transmission under threshold modulation

**DOI:** 10.1101/2020.12.13.422611

**Authors:** Wei-Mien M. Hsu, David B. Kastner, Stephen A. Baccus, Tatyana O. Sharpee

## Abstract

Modulation of neuronal thresholds is ubiquitous in the brain. Phenomena such as figure-ground segmentation, motion detection, stimulus anticipation and shifts in attention all involve changes in a neuron’s threshold based on signals from larger scales than its primary inputs. However, this modulation reduces the accuracy with which neurons can represent their primary inputs, creating a mystery as to why threshold modulation is so widespread in the brain. We find that modulation is less detrimental than other forms of neuronal variability and that its negative effects can be nearly completely eliminated if modulation is applied selectively to sparsely responding neurons in a circuit by inhibitory neurons. We verify these predictions in the retina where we find that inhibitory amacrine cells selectively deliver modulation signals to sparsely responding ganglion cell types. Our findings elucidate the central role that inhibitory neurons play in maximizing information transmission under modulation.

## 1. Introduction

The need to use efficient representations within the nervous system currently provides one of the leading frameworks for understanding neural computation. This framework accounts for a number of different properties of neural responses [1, 2, 3, 4, 5, 6, 7, 8, 9, 10, 11, 12, 13, 14, 15], including optimal ways for neural circuits to adapt to statistically consistent changes in the input statistics [1, 16, 17, 18]. However, it is also important to consider the case where information transmission occurs in the presence of fluctuations in input statistics that might not be strong enough, or persist for long enough time, to trigger full-scale adaptation. These types of fluctuations are nevertheless important to take into account because they can evoke and/or represent modulatory influences from other circuits, as is ubiquitous in the brain. For example, modulatory influences include contextual or top-down signals about input properties on the scales larger than that of the neuron’s primary receptive field, which closely follows the neuron’s linear or the so-called classical receptive field [19]. Such contextual effects underlie figure-ground segmentation, motion selectivity, motion reversal or anticipation and other predictive effects in the retina the retina [20, 21, 22]. These effects are also prominent in the cortex where they include cross-orientation suppression [23, 24] and other non-classical receptive field effects in visual [25, 19] and auditory [26] cortices. Threshold modulation can also result from the direct action of neuromodulatory circuits [27] that represent changes in arousal and attention [28, 29, 30]. The ubiquity of modulatory signals makes it essential to consider how they may influence the properties of maximally informative neural circuits.

It turns out that modulation has surprisingly non-trivial effects on information transmission. On one hand, for a sensory circuit, modulation of neuronal threshold that is independent of the primary sensory input is bound to decrease the information that this circuit can transmit about that primary input. On the other hand, we will show that modulation always decreases information less than an equivalent increase in the primary noise. We further show that the negative impacts of modulation can be nearly eliminated if it is directed to a subset of sparsely responding neurons in a coupled neural circuit. In this way, the neural circuit can take advantage of the flexibility afforded by modulation of its response properties without suffering a reduction in information transmission.

We test predictions of this theory on responses of pairs of retinal ganglion cells (RGCs) that encode the same temporal fluctuations of light intensities but with different thresholds [31]. These cells have been termed adapting and sensitizing based on their short term plasticity, but for the present analyses in steady-state conditions, the main differences between these cell types are that adapting cells have higher thresholds and larger noise levels than sensitizing cells. Previous maximally informative solutions for pairs of neurons accounted for many aspects of these neurons’ responses, including why these two separate cell types are observed among Off neurons but not among On neurons [14]. However, some noticeable quantitative differences between theory and experimental measurements were left unexplained [14]. Recent studies have pointed out that incorporating multiple noise sources could affect the predictions for threshold differences between cell types [15]. Therefore, we set out to determine whether modulatory effects on a cell’s threshold would influence the theoretical predictions, bringing them into better agreement with experimental measurements. After testing a number of scenarios, we found that a model where a secondary pathway modulated the threshold of the primary pathway for each cell type (Box 1) could quantitatively account for the measurements of threshold differences between cell types, across several different contrasts. We envision that this threshold modulation occurs even for a fixed contrast, and in the case of the retina derives from contextual modulation from inputs on scales larger than neuronal receptive field center, or for cortical neurons, the classical receptive field [32].

Fitting the maximally informative model with threshold modulation to the retinal data also made it possible to separate the observed neural variability into the contributions due to threshold modulation and noise in the primary pathway. We found that higher noise levels of adapting cells can be fully explained by larger threshold modulation experienced by these neurons compared to those experienced by sensitizing cells; the primary pathway noise levels were similar for both cell types. Mechanistically, threshold modulation in adapting cells could be implemented as additional input from inhibitory amacrine cells. To confirm this prediction, we then directly recorded from and manipulated amacrine cells. These experiments revealed a more reliable distance-dependent input from amacrine cells to adapting cells compared to sensitizing cells, consistent with the scheme where amacrine cells modulate the thresholds of adapting cells.

The theoretical results are obtained here using basic concepts of information theory. Therefore, they should apply not only in the retina, but also in the cortex and other neural circuits. The results highlight the importance of using inhibitory neurons to deliver modulatory signals into a circuit, which can provide a new framework for understanding the function of inhibitory neurons in the brain.

### Box 1

Two-pathway model of information transmission in the presence of threshold modulation

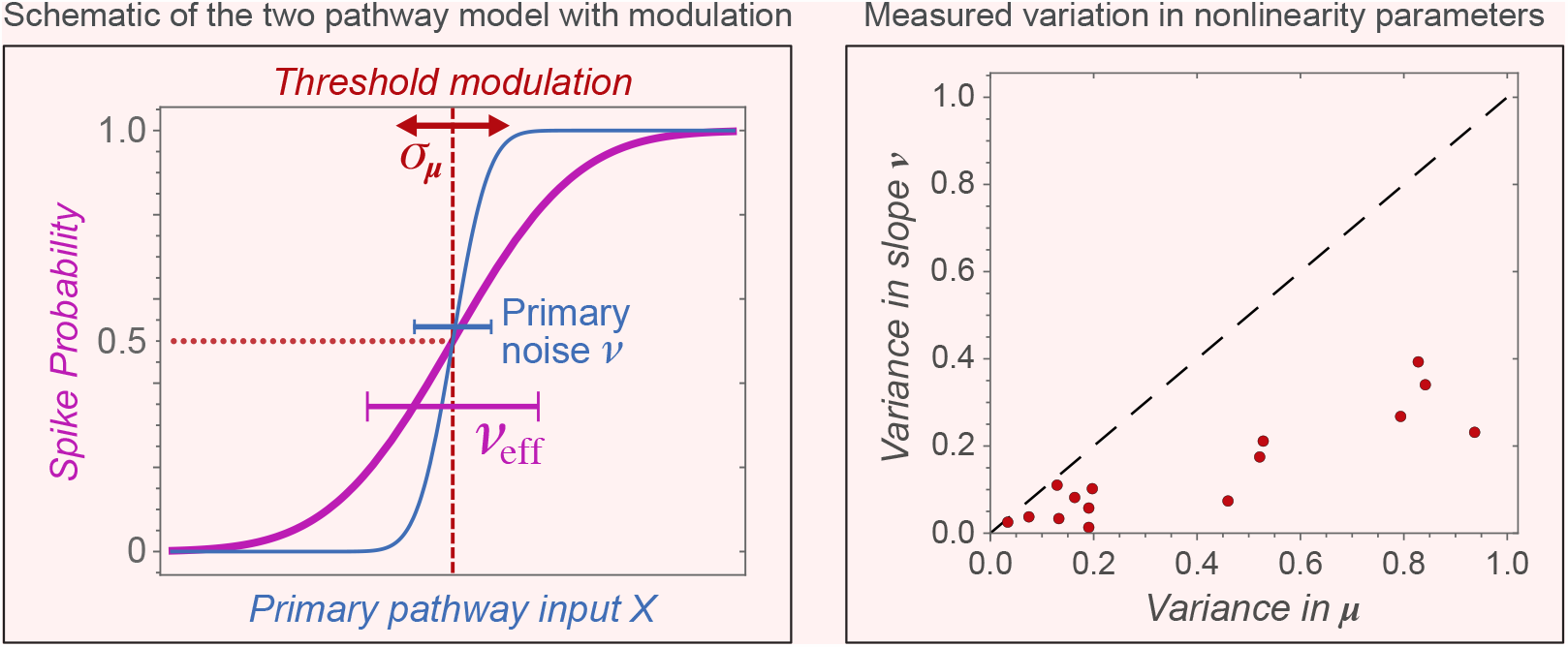

**The experimentally observed neural nonlinearity reflects two noise sources:**

- the intrinsic noise *ν* in the primary pathway
- threshold modulation that occurs on longer time scales with variance *σ*_*µ*_.
- Over time, the observed nonlinearity is an average over different threshold positions 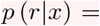 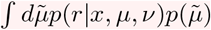 and has an effective width 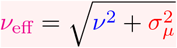

Right panel: Threshold variation over time is much stronger than variation in the primary noise.

The **mutual information** is computed in two steps:

1. On short time scales, mutual information between is computed for a fixed threshold *µ*:

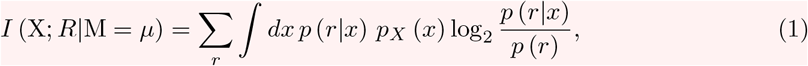

where *x* is the filtered stimulus according to the spatiotemporal receptive field of the neuron, and *r* ∈ {0, 1} represents the response of a single neuron before the incorporation of the modulation in the secondary pathway (*σ*_*µ*_ = 0, *ν*_eff_ = *ν*).
2. On longer time scales, we average the mutual information over the varying threshold value 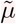:

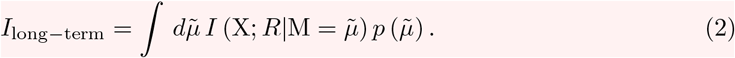

Here, 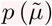 describes the distribution of threshold values.

The information in Eq. (2) is actually the so-called conditional mutual information [33] *I*(X; *R* M) between the input and the responses of the primary pathway, conditional of the signals *µ* from the modulatory pathway. As such, this information differs from the full information provided jointly by modulatory and primary pathways only by the term *I*(X; M): *I*(X; *R*|M) = *I*(X; {*R*, M}) − *I*(X; M), where *I*(X; M) represents information provided solely by the modulatory pathway. Because *I*(X; M) does not depend on the parameters of the nonlinearity of the primary pathway, it can be dropped when searching for the maximally informative properties of the primary pathway. Thus, one can find the maximally informative setting for the primary pathway and the optimal modulation by maximizing information from Eq. (2). These arguments generalize to the case of multiple neurons where one evaluates information between inputs X to the primary pathway of each neuron and the vector of responses across the neural population **R** = {*r*_*i*_}, *r*_*i*_ ∈ {0, 1}.

## 2 Results

### 2.1 Impact of threshold modulation on information transmission

To understand information transmission in the presence of threshold modulation, we modeled responses of individual neurons as binary, 1 or 0, corresponding to the presence or absence of a spike in a small time bin, respectively. Spiking probability is modeled as a threshold crossing event, with a threshold (*µ*) and a noise level (*ν*), which determines the variation in neural responses for a given input value. When parameter *ν* is small, there is only a small range of stimuli for which neuronal responses varies strongly from trial-to-trial with a probability ∼ 0.5. For inputs that are either much greater or smaller than the threshold *µ*, the spike probability is nearly certain, with values close to either 1 or 0, cf. Box 1. When the parameter *ν* is large, the range of stimuli with uncertain neuronal responses is large. The increase in the uncertainty in neural responses with *ν* can be quantified using a quantity known as noise entropy [34], which represents the average uncertainty in the neural responses across different stimuli.

This model of neural responses yields a saturating nonlinearity shown in Box 1 and described by the following equation:

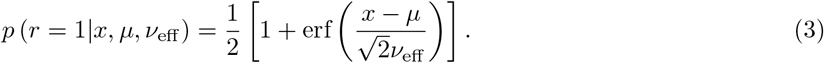

In this equation, we write *ν*_eff_ instead of *ν* to emphasize the fact that the observed noise in neural responses represents actually a joint effect of multiple different types of noise [15]. Here we will focus on two types of noise: the “primary” noise *ν* that arises in the direct afferent circuitry for each cell, and the secondary source of variability that arises from the modulation of the threshold *µ* of the primary pathway and acts on longer time scales. On short time scales, similar to those of the spike generating process, the threshold value does not vary, and variability in neural responses is described by *ν* only. On long time scales (∼ seconds), which are necessary to measure the neural input-output function, its width is described by

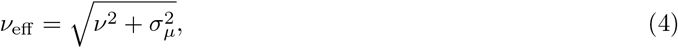

We note that, in principle, noise *ν* in the primary pathway can itself also be subject to modulation, not just the threshold *µ*. This modulation would also increase *ν*_eff_. However, in practice, we found that variation in *ν* was much weaker (Box 1, right panel). Therefore, in what follows, we focus on the effect of modulation on changes in the threshold.

To analyze the impact of threshold modulation on information transmission, we compute the mutual information in two steps, cf. Box 1. First, the mutual information is computed for a given value of the threshold, and then the results is integrated over threshold positions (Box 1). This procedures yields (up to a term that does not depend on parameters of neuronal nonlinearities) the mutual information provided about inputs by neural responses together with modulation signals.

We start by considering the impact of threshold modulation on single neurons. Here, modulation always decreases information transmission (Fig. 1A). However, for an equivalent amount of variance, modulation decreases information less than does primary noise. Therefore, if the system has a choice between reducing the primary noise or reducing modulation, it is always preferable to reduce the primary noise first, cf. Fig 1B.

**Figure 1:**
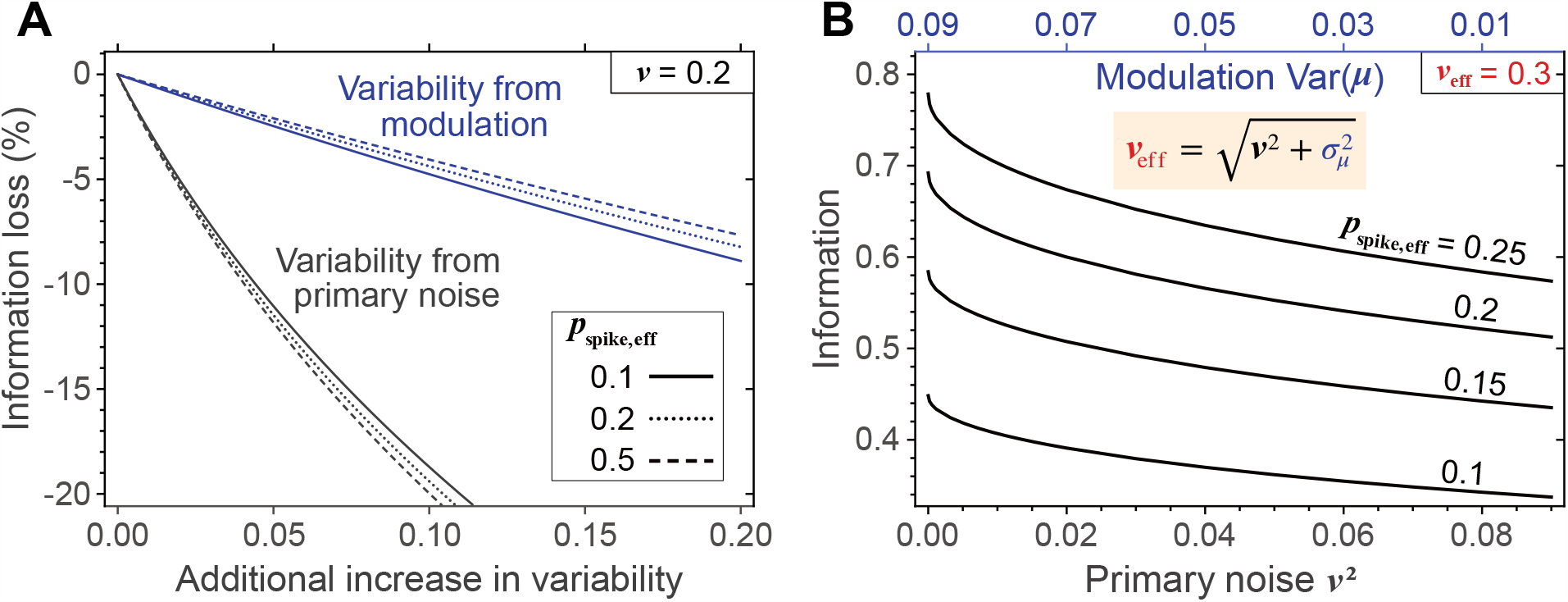
Impact of threshold modulation on information transmission. (**A**) The difference in information before and after adding different types of variability: either modulation (blue lines) or primary noise (black lines). Both of types of variability decrease information, but modulation (blue lines) decreases information much less than the primary noise (black lines). We note that both the primary noise and the modulation also increase the spike rate. Therefore the baseline information (without modulation) is computed for the higher rate that matches the rate in the presence of modulation. (**B**) The stronger detrimental effects of primary noise on information transmission compared with modulation are shown here for the case where primary noise and modulatory variance are constrained to sum 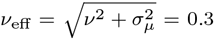. In this case, the smaller the primary noise (bottom *x*-axis), the larger the information (*y*-axis), despite the corresponding increases in modulatory variance (top *x*-axis).

The effect becomes more interesting in groups of neurons, starting with pairs of neurons. Here, we find that if modulation is directed to the neuron with the lowest firing rate in the group, then the negative effect of modulation is almost completely removed, cf. Fig. 2, panels A and B. In these calculations, the firing rates were assigned to maximize information while constraining the average spike rate across the neurons (Fig. S2). We find that one can apply much larger modulation to a single neuron than the modulation distributed to many neurons and still have less of a decrease in information. Selective application of modulation also maximized information in groups of three neurons (Fig. 2C,D). With three neurons, information was maximally preserved under modulation when it was applied to the neuron with the smallest spike rate. The most detrimental effects of modulation were observed when modulation was applied to the neuron with the largest spike rate. This was followed by progressively better results if modulation was applied equally to all neurons or to the neurons with the intermediate spiking rate. However, these intermediate cases still led to worse performances compared to the case where modulation is directed to the neuron with the lowest spike rate (Fig. 2D). The degree of protection from modulation-induced loss is higher for the three-neuron circuit compared with a two-neuron circuit (Fig. 2D). This suggests that the benefits of including a sparsely responding neurons can be larger in large groups of neurons.

**Figure 2:**
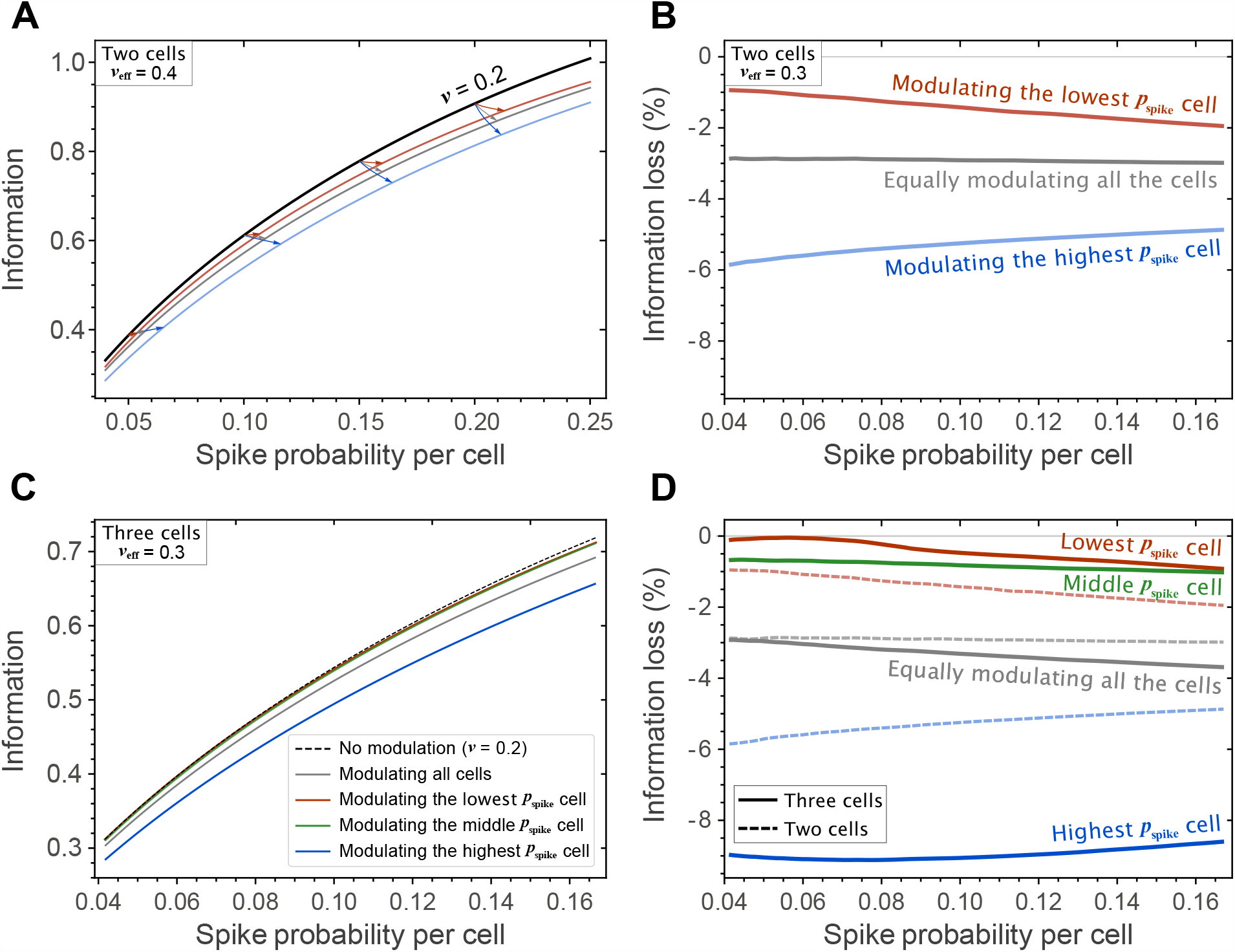
Modulation directed to sparsely responding neurons protects against information loss in the presence of modulation. (**A**) The information loss is smallest when only the lowest-spiking neuron (red line) receives modulation, compared to modulating all neurons (gray line) or the highest-spiking one (blue line). Black line shows information in the absence of modulation. The primary noise *ν* = 0.2 for all cases, lines with modulation have the same averaged effective noise *ν*_eff_ = 0.4 after modulation. Arrows describe how points on the unmodulated curve change in terms of information and spike rate upon adding the same amount of overall modulation. The red and blue arrows have different final values for spike rate because the modulation-induced increase in the spike rate depends on the initial spike rate values and is different for the lowest and highest spiking neuron in the pair. The averaged effective noises after modulation are *ν*_eff_ = 0.3 for all curves. The spike rates were optimized to yield maximal information for a given average spike rate. The corresponding rates are shown in Fig. S2. Same as (**A**) but in terms of percentage of information loss to show the results on an expanded scale. (**C**,**D**) Same as **A** and **B** but for three neurons. In (**D**) results from (**B**) pertaining to pairs of neurons are re-plotted using dashed lines for comparison. Green lines shows the case where modulation is directed to the neurons with intermediate spike rates, other colors are the same as for pairs of neurons. Directing modulation to the most sparse neurons yields the smallest information loss from modulation. Modulation can be more fully compensated in three-neuron groups compared to two neurons, for smaller spike rates.

We also examined the case where neurons have the same thresholds and spike rates, as can be optimal for high values of the primary noise [14]. In this case we found that the optimal ways to apply modulation differed depending on whether same-threshold neurons had small or large spike rates, cf. Fig. S3. In the case where neurons had small rates, it was optimal to apply modulation equally to both of them. In the case where neurons had large response rates, it was optimal to direct modulation to one of the neurons than split it equally to both neurons (Fig. S3). The application of modulation lowered the spike rate in the target neurons. The implication from these results therefore is that if a large neural circuit contains neurons of the same type with small spike rates, such as for example the adapting cells in the retina, then modulation should be applied selectively to the class of neurons with sparse responses and equally within this class neurons.

Why is it beneficial to direct modulation to the neuron with the lowest spike rate? An intuitive explanation for this phenomenon can be obtained by considering the shape of the information function for a single neuron with respect to its threshold (Fig. 3A). This function is concave for small thresholds and convex for large thresholds. This is important because concave functions decrease their value upon averaging of their inputs, as occurs as a result of threshold modulation, while convex functions increase their value. This means that neurons with small thresholds, i.e. high spike rates, will suffer a decrease in information transmission upon modulation, cf. Fig. 3B. In contrast, neurons with large thresholds, i.e. small spike rates, will increase information transmission upon threshold modulation. The lower the spike rate, the greater is the increase in the information transmission with modulation. This explains why directing modulation to the neuron with the lowest firing rate is more beneficial than directing modulation to neurons with higher firing rate. As a related points, one can also notice in Fig. 2B that the protection against modulation-induced loss in information transmission decreases with the average spike rate.

**Figure 3:**
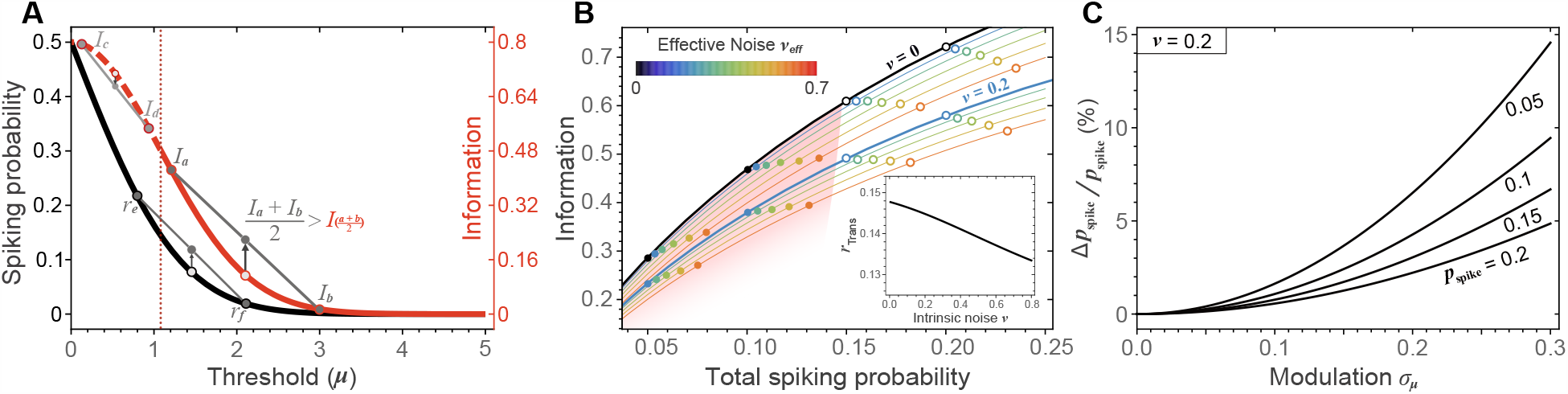
Modulation induced transition in information transmitted as a function of spike rate. (**A**) Spike probability is a convex function of threshold position (black line). In contrast, information (red line) changes convexity as a function of threshold. When a function has positive convexity (solid segments of the curve) the average of its two values at points *a* and *b* is always larger than the function value at (*a* + *b*)*/*2. In this regime, fluctuations increase information transmission. The opposite is true for regions of negative convexity (dashed-curve). As a result, fluctuations in threshold decrease information when thresholds are low and increase information when threshold are high, i.e. when neurons respond sparsely (**B**) Threshold modulation increases information transmission when spike rates are small (filled dots) but decreases it when spike rates exceed a certain transitional value (open dots).Shaded pink region denotes the value where modulation increases information transmission. Thick solid lines show information in the absence of threshold modulation 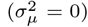, for two noise levels *ν*_1,2_ = 0 (black) and 0.2 (light-blue). Thin solid lines and the eight series of color-dots on them show how curves shift upon introduction of threshold modulation. Each series of color-dots evolves from the same intrinsic noise (*ν*) and threshold (*µ*). Color denotes the resulting effective noise 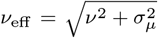. (Inset) The transitional value in response rate is plotted as a function of the intrinsic noise. (c) Modulation increases response rate.

At this point, it is important to clarify that this increase in information transmission with modulation is accompanied by an increase in the spike rate. Unlike information, the firing rate function is convex for all values of its argument (Fig. 3A). As a result, modulation always increase the spike rate (Fig. 3C). The increase in the information from modulation is less than it would have been if the rate was simply increased by lowering the threshold, without the modulation. As a result, the information vs. rate curve in the presence of modulation has the same shape as in the absence of modulation, just with reduced information for a given rate.Thus, these results are consistent with those in Fig. 1A showing modulation decreases information. It is just that the increase in information upon modulation can nearly completely match the increase that would have been observed if the firing rate was increased without modulation.

The conclusions from the theoretical analyses of information transmission in the presence of threshold modulation indicate that modulation should not be equally distributed to all neurons in the target circuit. Instead, it should be directed to the neuron with the lowest spike rate with inhibitory signals. The use of inhibitory signals ensures that the rank-ordering of neurons does not change under modulation, and the neuron that receives modulation does not get its spike rate raised. They also illustrate the need to use neurons with diverse spike rates, because the average spike rate in the circuit sets the upper limit on the amount of information that this group of neurons can transmit, with or without modulation. To have the capability to transmit large amounts of information, the circuit has to include neurons with large spike rates. Including neurons with small response rates and directing modulation to them helps maintain information transmission near its maximal levels in the presence of modulation.

### 2.2. Retinal input-output functions are maximally informative under threshold modulation

We now test how these predictions using responses of pairs of cells in the retina that differ in their average spike rates. The adapting and sensitizing cells are two cell types that represent the same temporal pattern of light intensity modulation but have different thresholds. Our first analysis is to fit the maximally informative model with modulation to the responses of pairs of adapting/sensitizing cells. The fit was made while requiring that the effective noise and the average spike rate for the pair matched experimental measurements (see Methods for details). The fit yields estimates for threshold modulation and primary noise for each neuron in the pair as well as an estimate for the difference in their thresholds. These estimates can then be compared to direct experimental measurements of these variables.

We find that the inferred amount of noise in the primary pathway was similar for both adapting and sensitizing cells (Fig. 4A). However, the threshold modulation was substantial for adapting cells and very close to zero for the sensitizing cells (Fig. 4A). The fitting results were consistent across cell pairs. (Table S1). Thus, the differences in the effective noise that are observed between these two cell types [31] are due to differences in threshold modulation. We also note that threshold modulation was small in sensitizing cell even relatively to their thresholds (the modulation was ∼ 100 times smaller for sensitizing cells compared to adapting cells, whereas their thresholds are only approximately half as small as those of adapting cells).

**Figure 4:**
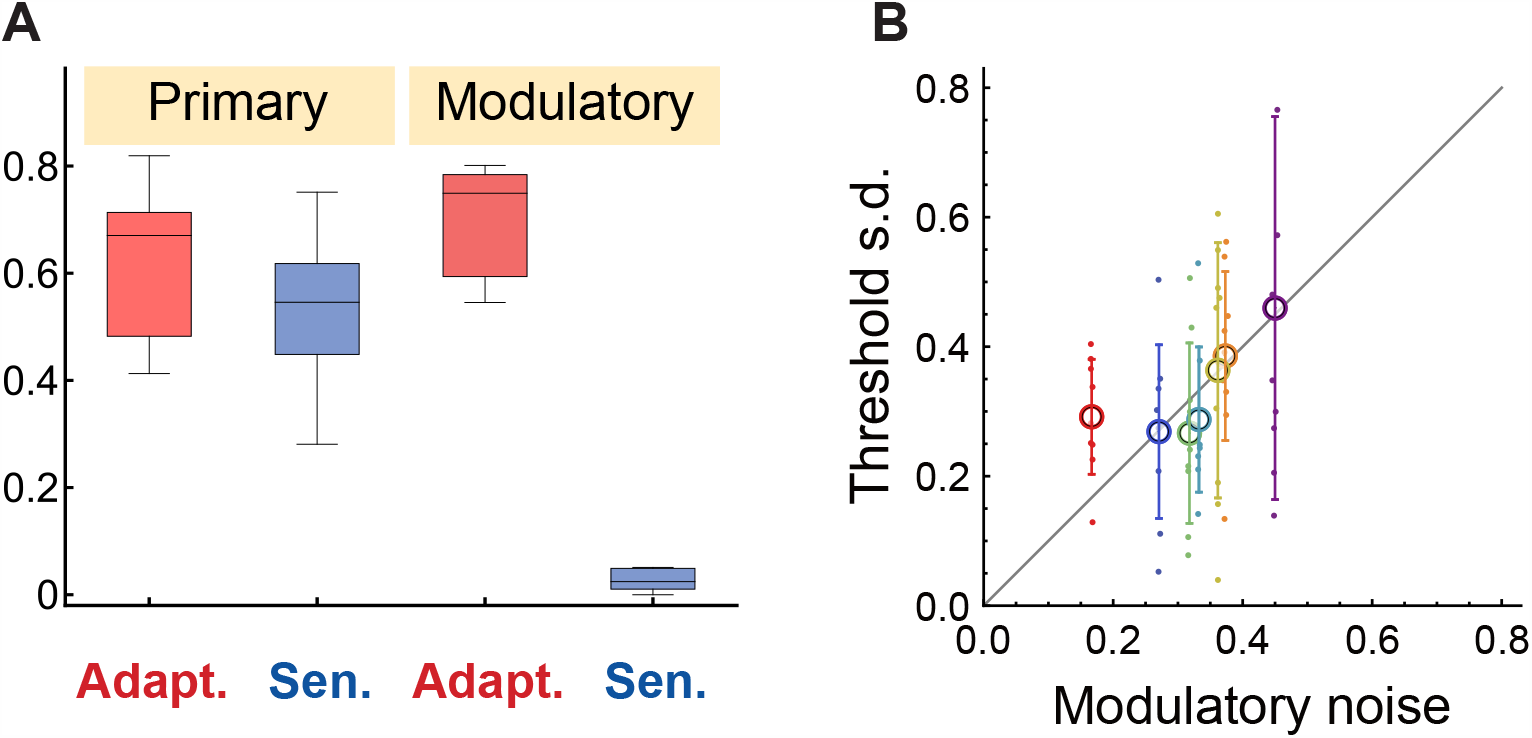
Experimentally observed threshold variation matches maximally informative values. (**A**) Intrinsic neural noise and threshold modulation inferred using the maximally informative model with modulation from retinal data. Both neural types have comparable amounts of intrinsic neural noise (*ν*_*i*_) but distinct levels of threshold modulation (*σ*_*µ,i*_). All noise types varied linearly with the stimulus contrast, except for modulatory noise in the sensitizing cells, which was small and contrast-independent. (**B**) The experimentally observed threshold variation across adapting cells is positively correlated with threshold modulation inferred from the maximally informative model (*r* = 0.3, *p* = 0.015). Both axes are in units of contrast. Colors denote different neurons. Data points for the same neuron/color represent measurements from different input contrasts.

The threshold modulation values predicted by the maximally informative model with modulation can be compared with direct experimental estimates of their threshold modulation. To compute the amount of threshold modulation that is observed experimentally, we estimated neuronal nonlinearities from shorter data sub-sets (1/4 to 1/6 compared to the full dataset). Each nonlinearity was fit with a logistic function to determine its threshold value. We find that the observed variation in thresholds for a given adapting cell matches those estimated using the maximally informative model (Fig. 4B, paired non-parametric t-test *p* = 0.73). [This analysis was only carried out for adapting cells, because threshold modulation was negligible in sensitizing cells]. Those adapting cells that had larger variance in thresholds across trials also had larger values of threshold modulation as indicated by fitting the maximally informative model to the full set of their response (the correlation was statistically significant, with *p* = 0.015, Fig. 4B). These analyses add credence to the use of the maximally informative model with modulation as a method for separating the noise component that is due to threshold modulation. They also indicate that the observed threshold modulation in adapting cells is maximally informative given their other parameters, such as the primary noise and firing rate.

Another prediction that one can obtain from the maximally informative model with modulation pertains to the differences in the thresholds between adapting and sensitizing cells. Previous predictions for the threshold differences obtained for pairs of neurons without taking modulation into account yielded values that were systematically larger than those observed experimentally [14], replotted in Fig. 5 with black line. We find that the maximally informative model with modulation provided more accurate predictions for thresholds differences between pairs of neurons than the model with no modulation, cf. Fig. 5. Statistically, the threshold difference (in units of contrast) between adapting and sensitizing cells were consistent between the average values across contrasts for each cell pairs from the maximally informative model and experimental measurements (paired non-parametric t-test *p* = 0.14). By comparison, the model with no modulation yielded systematically greater threshold differences that is observed experimentally (black line in Fig. 5). We note that experimental data points show larger residual variation across different contrasts than our model indicates. The reason for this is that, in the model, noise components and threshold modulation for adapting cells were constrained to change linearly with contrast (to reduce the number of fitted parameters, see Methods). Thus, the model was not meant to predict residual variation across contrasts that remains after rescaling inputs by their contrast. Other than this variability, the predictions of the maximally informative model with modulation for threshold differences between adapting and sensitizing cells are fully consistent with experimental measurements (*p >* 0.14, Fig. 5B).

**Figure 5:**
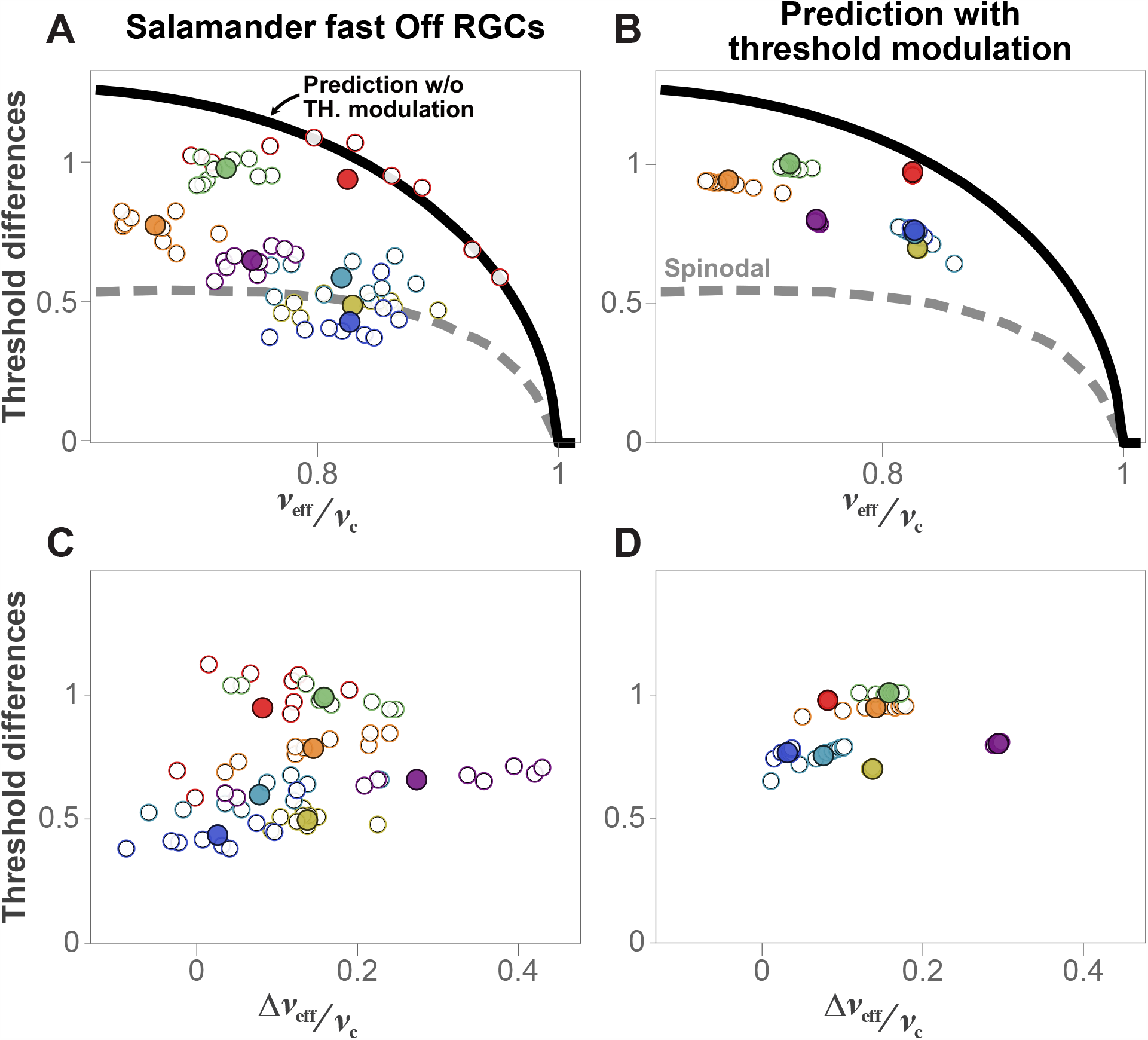
Maximally informative model with modulation accounts for threshold differences between adapting and sensitizing cells. Threshold difference between adapting and sensitizing cells is plotted as a function of average (top row) and difference (bottom row) in the effective noise between the two neurons. Columns show data (left), maximally informative predictions with modulation (right). Different colors denote different cell pairs. Open circles represent data for a given contrast, filled circles show the average across contrasts. Black lines show predictions for threshold differences without threshold modulation. Gray dashed lines denote spinodal lines that separate regions where information has two maxima vs. a single maximum. Points close to the spinodal lines (e.g. blue, light blue, and light green) are more difficult to fit because they mark the region where one of the maxima ceases to exist. This pushes the interpolated solutions away from the spinodal line (c.f. Fig. S1). Despite these technical issues, the overall distribution of mean threshold values normalized across contrasts was not statistically different between fitted and experimental values, *p* = 0.14.

### 2.3. Amacrine cells as a source of threshold modulation for adapting cells

One of the key predictions of the theory is that modulation should be directed to neurons with low spike rates. However, as we have seen above, modulation increases the spike rate (Fig. 3C), albeit by moderate amounts. One way to minimize the risk of altering the rank-ordering of neurons in terms of their spike rate is to deliver it with inhibitory neurons. In this way the neuron that is undergoing modulation will automatically have its threshold raised and spike rate lowered. This is consistent with our observations in the retina where adapting neurons, which undergo modulation, also have larger thresholds and smaller spike rates. In the retina, inhibitory amacrine cells could be the source of that input (Fig. 6A). If amacrine cells provide stronger inputs to adapting cells than the sensitizing cells, then this would simultaneously explain why the thresholds of adapting cells are higher and more variable than those of sensitizing cells. The fact that both the mean threshold and its modulation varies approximately linearly with contrast is also consistent with this wiring scheme. Inputs to and from amacrine cells just need to be scaled by contrast just like inputs within the primary pathway for the adapting and sensitizing cells.

**Figure 6:**
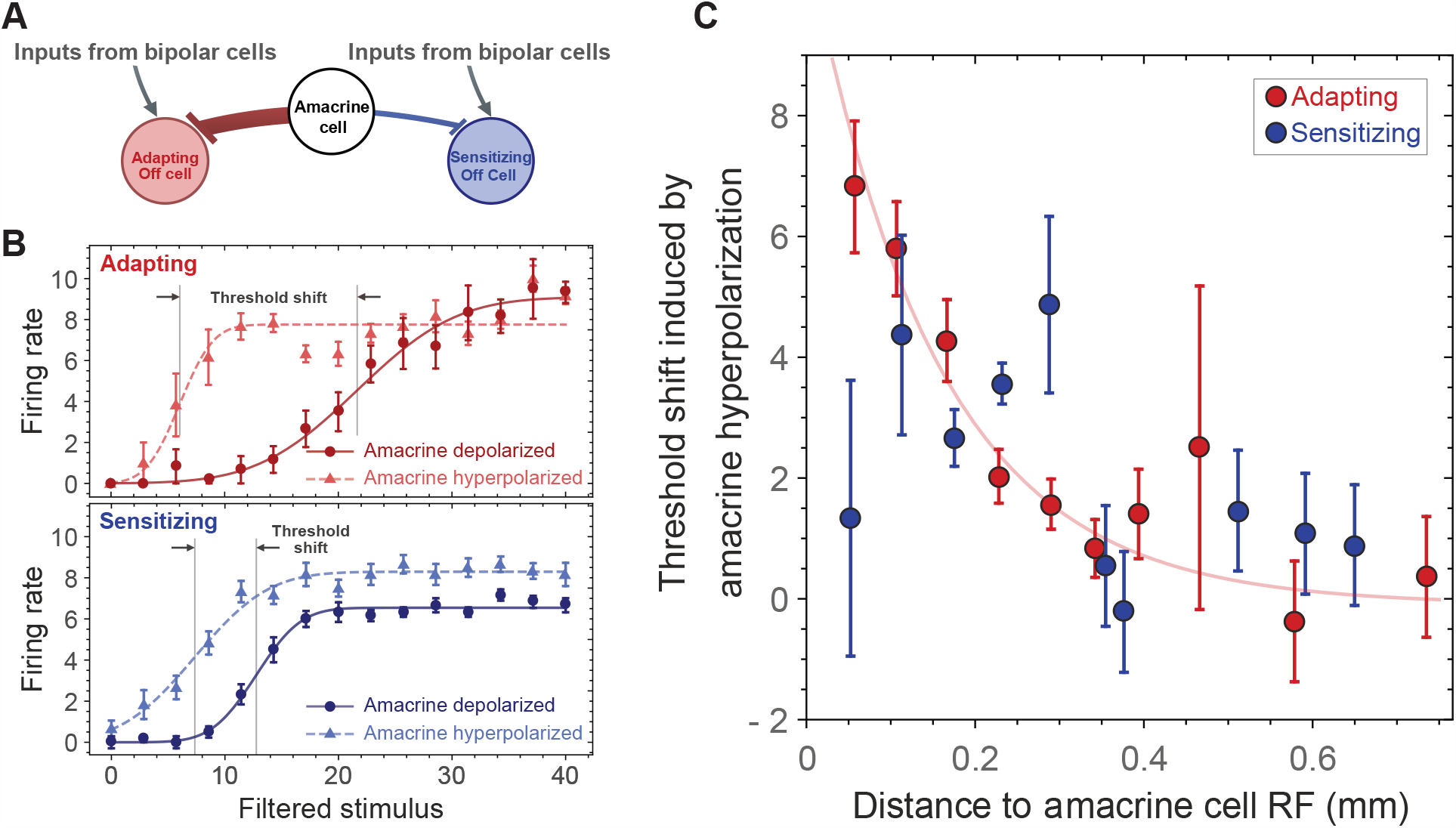
Distance dependent inputs from amacrine to adapting cells. (**A**) Inferred model of the presynaptic circuitry of the two types of Off retinal ganglion cells based on observed differences in the strength of the modulatory pathway. (**B**) The nonlinearity of Off ganglion cells during the depolarizing (dot) and hyperpolarizing (triangle) current injection into the amacrine cell. The solid and dashed curves show the fit with sigmoid function. cells. The distance between the receptive field (RF) of the amacrine cell to that of the adapting cell was 0.090 mm, 0.101 mm to the RF of the sensitizing cell. (**C**) The amount of inhibitory input from amacrine cells to the adapting cell decreases with distance significantly (*p ×* 10^*−*8^, f-test). [Inhibition may be direct or polysynaptic, through circuitry involving bipolar cells or other amacrine cells.] The dependence on distance was not statistically significant for sensitizing cells (*p* = 0.9). Solid lines show the exponential fits with distance.

We tested this hypothesis by performing a separate set of experiments to analyze how the hyperpolarization and depolarization of sustained Off-type amacrine cells by intracellular current injection affected responses of nearby adapting and sensitizing cells recorded simultaneously with a multielectrode array (see Methods and Fig. 6). Specifically, we analyzed the change in the mean threshold of adapting/sensitizing neurons between hyperpolarization and depolarization of the amacrine cell. When an amacrine cell is hyperpolarized (depolarized), this decreases (increases) its inhibition onto neurons it is directly connected to. Although we do not assume that there are direct connections between amacrine cells and the ganglion cells we recorded (the connection could be polysynaptic, through circuitry involving bipolar or other amacrine cells), this approach measures the functional effect of individual amacrine cells. In Fig. 6C we plot the change in the threshold as a function of distance between the receptive fields (RFs) of the amacrine cell (that was subjected to hyperpolarization/depolarization) and the adapting/sensitizing cell whose nonlinearity was measured to estimate its threshold. In the case of adapting cells, there was a clear and statistically significant dependence of the amount of threshold shift as a function of the distance to the amacrine cell RF center (*p* = 8 × 10^*−*5^ F-test compared with null hypothesis of no dependence on distance). The dependence was not statistically significant in the case of sensitizing cells (*p* = 0.9). Thus, these data support the hypothesis that the larger thresholds of adapting ganglion cells arise as a result of inhibition from the amacrine cells, and that this inhibition also brings with itself stronger threshold modulation.

## 3. Conclusion

In this work we analyzed information transmission in the presence of threshold modulation. There are two main conclusions. The first conclusion is that modulation should not be equally applied to all neurons in the circuit. Instead it should be directed to select neurons, preferably those with the low spike rates in the circuit. The second conclusion describes the central role that inhibitory neurons play in delivering modulatory signals into the circuit. These conclusions are obtained by through basic analyses using information theory, and therefore should apply to all neural circuits. We now discuss the implications of these conclusions, with a focus on cortical circuits.

The first conclusion highlights the need to form circuits using neurons with different spike rates. The large number of sparsely firing neurons in the cortex have long presented a puzzling observation [35]. The chief explanation offered so far is that sparse response reflect due to metabolic constraints. However, one could have hypothetically used a smaller number of neurons with higher spike rates, if metabolic constraints were the leading cause for the sparseness of neural responses. The information-theoretic analyses in the presence of modulation offer a different explanation. Neural circuits need to have neurons with both high and low firing rates in order to transmit large amounts of information in the presence of modulation. High firing neurons make it possible to transmit large amount of information, whereas neurons with small spike rates protect against loss of information transmission in the presence of modulation.

The second conclusion describes a rather unexpected role for inhibitory neurons as intermediaries for delivering modulation signals. This set up helps to ensure that low-spiking neurons that receive modulation remain in this regime under varying modulation levels. We find support for this prediction in the retina where inhibitory amacrine cells send modulatory signals to sparsely spiking adapting cells. If modulation were delivered to adapting cells via excitatory pathway, then this would risk making their spike rate greater than that of sensitizing cells and losing protection against negative effects of threshold modulation on information transmission.

The same role appears very plausible for inhibitory neurons in the cortex. There, inhibitory neurons expressing the vasoactive intestinal peptide are the major target of neuromodulatory inputs as well as modulatory, context-dependent inputs from higher-order cortical areas [36]. Similarly, somatostatin expressing inhibitory neurons use this neuropeptide as a co-transmitter with GABA to modulate the activity of local neurons [37]. The slow action of neuro-peptides, such as somatostatin, conforms with our modeling framework where modulation changes neuronal threshold on slower time scales than those on which the primary activation pathway operates. We note also that all of the other inhibitory neurons, including parvalbumin-positive inhibitory neurons, are directly responsive to neuromodulators such as acetylcholine and serotonin [38]. Furthermore, even when neuromodulators, such as acetylcholine, act directly on excitatory neurons, they exert first an inhibitory response [39] in their target neurons. The information-theoretic results offer a new explanation for this tight link between neuromodulatory and inhibitory circuits in the brain.

## Methods

### Experimental preparation

We use a combination of new and previously published experimental data [31]. Full details of the experimental procedures for measuring neural nonlinearities are provided in [31]. Briefly, uniform field stimuli were drawn from a Gaussian distribution with constant mean intensity, *M*, of 10 mW/m^2^. Contrast is defined as *σ* = *W/M*, where *W* is the SD of the intensity distribution. Neurons were probed with flashes of nine different contrast values from 12% to 36% in 3% intervals. The contrasts were randomly interleaved and repeated. Each contrast was presented, in total, for ≥ 600 s. For the calculation of the response functions, the first 10 s of data in each contrast were not used to allow for a better estimation of the steady state.

### Intracellular recording

Simultaneous intracellular and multielectrode recordings from the isolated intact salamander retina were performed as described [40]. Sustained amacrine cells were distinguished from horizontal cells by their flash response and their spatiotemporal receptive fields, with horizontal cells lacking an inhibitory surround and being greater than 300 *µ*m in diameter. For the intracellular recordings the stimulus comprised of randomly drawn contrasts with contrast amplitudes that ranges from 0 to 40% Michelson contrast units, where Michelson contrast is defined as (*I*_max_ − *I*_min_) */* (*I*_max_ + *I*_min_). The flash amplitude varied randomly every 400 ms, the first 100 ms the flash was greater than the mean, from 100 to 200 ms the flash was lower than the mean, and for the last 200 ms the flash was at the mean luminance level. Changing the distribution of amplitudes slower than the integration time of ganglion cells allowed for a rapid measurement of the ganglion cell response function without having to also measure the ganglion cell temporal filter [41]. Synchronized to the visual stimulus, we injected from 100 to 300 ms, randomly interleaved, hyperpolarizing (−500 pA) or depolarizing (+500 pA) current pulses into the amacrine cell. The ganglion cell response function was calculated at the firing rate of the ganglion cell from 100 to 400 ms of each contrast amplitude. This focused on the off response of the ganglion cell.

### Maximally informative modulation model for two neurons

We begin by reviewing the main features of maximally informative solutions for two neurons obtained in the absence of threshold modulation [14, 42]. The most prominent feature of the mutual information is a bifurcation that occurs when noise decreases below a certain, critical value (Fig. S1). In the case where both neurons have the same noise levels *ν*_1_ = *ν*_2_, a single peak at zero threshold difference splits into two symmetric peaks upon decreasing noise level. Each of these peaks represents equivalent solutions obtained by exchanging neuronal indices. One of the peaks describes the case where *µ*_1_ *> µ*_2_ whereas the other describes the case where *µ*_1_ *< µ*_2_. When neurons have different noise values *ν*_1_ and *ν*_2_, the peak with *µ*_1_ *< µ*_2_ becomes suboptimal if *ν*_1_ *> ν*_2_. Thus, the lower threshold neurons should have lower noise. This agrees with the intuition that a neuron which is more sensitive to small input fluctuations should have smaller noise. From the measurements of the average spike rate for the two neurons, one can predict the critical noise value below which one can expect to find neurons with different thresholds encoding the same filtered stimulus *x*. The critical noise value was indeed above the measured noise values for the adapting and sensitizing retinal ganglion cells (RGCs) [14]. In addition, one can make detailed predictions for the expected value *µ*_1_ − *µ*_2_ based on the measurements of other parameters *ν*_1_, *ν*_2_ and *µ*_1_ + *µ*_2_. Here, theoretical predictions were in qualitative agreement with experimental measurements, but quantitatively the observed threshold differences between the adapting/sensitizing neuron pairs were systematically smaller than those predicted based on maximizing information (Fig. 5A). We now show that taking into account threshold modulation brings theoretical predictions into agreement with experimental data.

To understand how threshold modulation affects maximally informative threshold positions, one may note that threshold modulation effectively smooths the information surface computed over long time scales (Fig. S4). In the regime where the mutual information has two maxima, it has the effect of bringing the maxima closer to each other. Another effect that proved necessary to take into account is that noise in the primary pathway can be larger for the neuron that experiences smaller threshold modulation, leading to a smaller overall effective noise value for that neuron. In this case, the information transmitted matches the smaller (local) of the two maxima of the information. In other words, the model allows for the possibility that coordination of neural thresholds between neurons might not be able to keep up with changes in input statistics for the circuit to match the properties of the global maximum of information. Instead, we observed that in some cases neural response properties match a local maximum of the information that required smaller adjustments in thresholds following the change in input statistics.

Taking both of these effects – threshold modulation and the possibility of local optimality – made it possible to account for the observed threshold differences between sensitizing and adapting cells. Each cell pair was probed with flashes of nine different contrasts, producing four experimental parameters of the neuronal nonlinearity (*ν*_eff,1_, *ν*_eff,2_, *µ*_1_, *µ*_2_) at each contrast. The maximally informative model also has six parameters (*µ*_1_, *µ*_2_, *ν*_1_, *ν*_2_, *σ*_*µ*,1_, *σ*_*µ*,2_). It can predict the difference *µ*_1_ − *µ*_2_ given a set of values for *µ*_1_ + *µ*_2_, *ν*_1_, *ν*_2_, *σ*_*µ*,1_, *σ*_*µ*,2_; only three of these five parameters are constrained by the measured input-output functions. Thus, the model is underconstrained for one value of contrast. However, experiments indicate that once neurons are adapted to a given value of contrast, parameters of experimentally measured nonlinearities increase approximately as a linear function of contrast [13, 31, 18, 43, 44]. We use this observation to fit the maximally informative model across contrasts. The resulting model has eight parameters altogether: the linear and offset terms with respect to contrast for each of the four noise terms (*ν*_1_, *ν*_2_, *σ*_*µ*,1_, *σ*_*µ*,2_). Because position of information maxima are affected by changes in any of these parameters, the maximally informative model can therefore be used to predict 27 independent measurements across contrasts (three values of *µ*_1_ − *µ*_2_, *ν*_eff,1_, and *ν*_eff,2_ for each contrast).

### Least-squared-fitting for parameters of the threshold modulation model from RGCs data

Base on the maximally informative modulation model, at a given (*µ*_1_ + *µ*_2_) the solution to threshold difference between a pair of adapting and sensitizing cell, ∆*µ*_model_, is nonlinearly dependent on the magnitude of each noise source (*ν*_*i*_, *σ*_*µ,i*_). This allows us to separately estimate the magnitude of these noise components from the neural data.

The results of least-square fitting were also constrained to match the observed values for *ν*_eff,*i*_. Seven pairs of adapting (index 1) and sensitizing cells (index 2) were probed by the nine different full range of contrasts (*σ* = 12% to 36% in 3% intervals [31]). The adaptive dynamics of noise level has been experimentally observed in many sensory systems[13, 31, 18, 43, 44]. Typically, the width of the transition region of the nonlinearity changes linearly with stimulus contrast (standard deviation). This adaptive process serves to optimize the information processing[18]. Here, we assume that both the primary (*ν*_*i*_) and the secondary (*σ*_*µ,i*_) noise sources are approximately linearly dependent on contrast (*σ*),

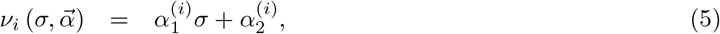

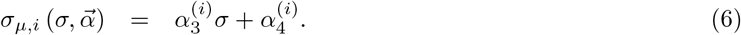

The effective noise also depends on contrast,

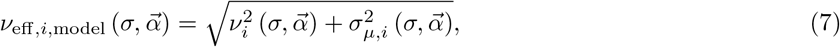

where *i* = 1, 2 denotes adapting or sensitizing neuron, respectively. The parameters 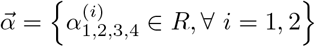 are to obtained by the least-squared-fitting for each cell pair while requiring them to also be consistent with *ν*_eff,*i*_ measurements from the shape of the nonlinearity. This model has eight parameters. Although formally it can be fit to data points for each individual cell pair, we reduced the number of parameters in half by focusing on the dominant term between the linear and contrast-independent terms for each type of noise. Initial fitting of the model indicated very small values for 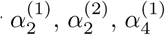 and 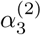. The final fitting reported here was obtained by setting these terms to zero, i.e., that noise in the primary pathway scales linearly with contrast for both types of cells; threshold modulation was set to be linearly increasing with contrast for adapting cells and to be contrast-independent for sensitizing cells.

The observed nonlinearities for a pair of adapting (index 1) and sensitizing cells (index 2) determine the threshold separations (∆*µ* = *µ*_1_ − *µ*_2_) and the effective noise levels (*ν*_eff,1 or 2_). For each cell pair, we aim to dissect two contributions to their *ν*_eff,1(or 2_): the one from the intrinsic noise level (*ν*) and that due to threshold modulation (*σ*_*µ*_), via minimizing the squared-error between the retinal data and the model predictions across the nine contrasts (*σ* = 12% to 36% in 3% intervals). Given a contrast (*σ*) a data point of a cell pair, 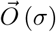, consists of three components,

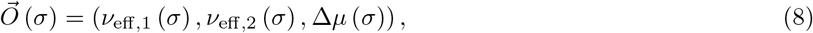

and so does our model 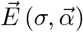,

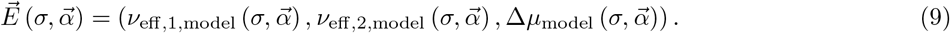

Here, 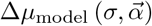 is the predicted threshold separation from our model, dependent on the intrinsic *ν*_*i*_ and modulatory noise *σ*_*µ,i*_ of each cell types,

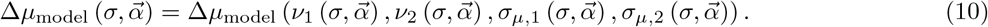

The predicted threshold differences (∆*µ*_model_) were firstly computed discretely in the grid space (*ν*_1_, *ν*_2_, *σ*_*µ*,1_, *σ*_*µ*,2_) and interpolated with Mathematica build-in function to construct the solutions between the grids. To avoid biasing the result by the component with largest error-bar, we standardize the 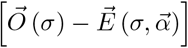 of each dimension with the inverse of its standard deviation. That is, the rescaling factors (weights) were

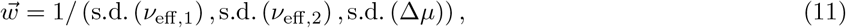

or more specifically,

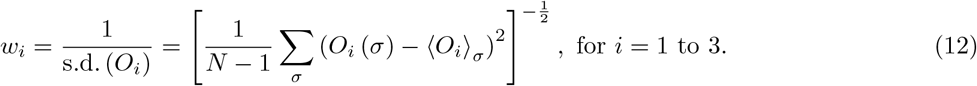

We defined the sum of weighted squared errors (or residuals) as

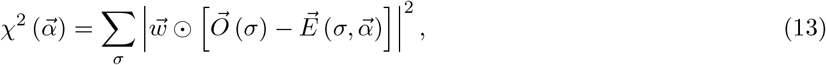

Where ⊙ denotes component-wise multiplication. The parameter 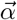 is the best-fit minimizing the weighted least-squared-error,

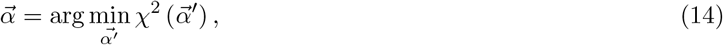

which predicts how the intrinsic (*ν*_*i*_) and the modulation noise (*σ*_*µ,i*_) depend on the stimulus contrast (*σ*). To quantify the goodness of fit, we use the variance (or reduced *χ*^2^)

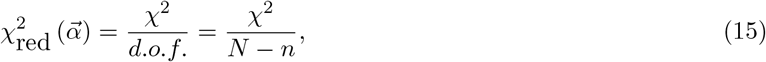

where *d.o.f*. = the number of degrees of freedom = *N* − *n*; *N* is the number of observations (nine contrasts in our case), and *n* is the number of fitted parameters. Note that by considering the threshold modulation, the predictions for the minimal threshold differences between the two cell types cannot go below the spinodal line. This makes it difficult to fit the data points adjacent to or below the spinodal region with our model. Therefore, the fitting results for three cell pairs did not adequately capture the trends (Fig. 5).

Finally, we also fit a single model across all cell pairs and contrasts. The resulting parameters (provided in the last row of Table S1) were consistent with average values of parameters fitted to individual cell pairs (Fig. 4).

### Computation of the information-theoretic limit on the response rate in the presence of threshold modulation

In the presence of small threshold modulation the effective information, Eq (2) can be approximated to the first order in threshold variance 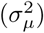 as:

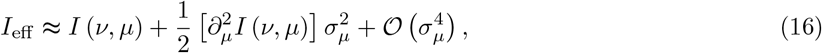

where the second term gives a good estimate on the information increment. The transition threshold value (*µ*_Trans*−*single_) is computed by solving the following equation:

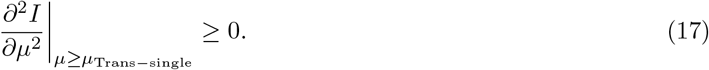

The corresponding transition values for the response rate and information are given by:

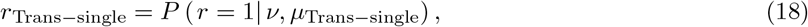

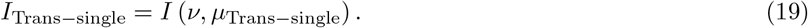

In the case of a pair of neurons, the effective information is approximated by:

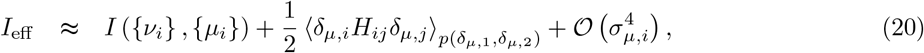

The transition values of thresholds (*µ*_Trans*−*pair,*i*_) are found from the following equation:

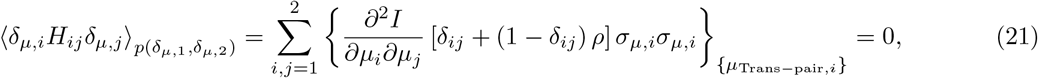

where 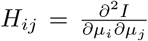 is the Hessian matrix of information *I*, and the probability distribution of threshold modulation *p* (*δ*_*µ*,1_, *δ*_*µ*,2_) is the multivariate Gaussian with correlation *ρ*. According to the estimated modulation from retinal data (Fig. 4 and Table S1), the sensitizing cells receive minimal threshold modulation. Therefore, we revise Eq. (21) to

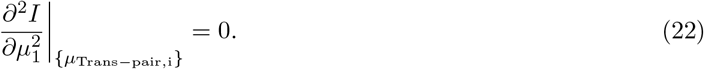

The above equation has multiple solutions corresponding to different linear combinations of the two thresholds {*µ*_Trans,*i*_}. We find the unique solution by constraining the difference in thresholds to match that in the data: *µ*_Trans,1_ − *µ*_Trans,2_ = ∆*µ*_data_. Finally, we can calculate the corresponding transition values for the response rate and information:

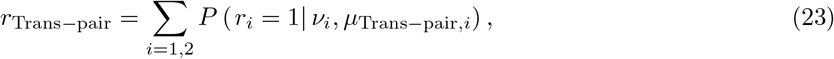

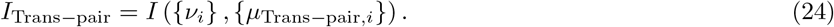

### Analysis of inhibition from amacrine cells versus RFs distance

To quantify the amount of inhibition from the amacrine cells to a ganglion adapting/sensitizing cells (Fig. 6), we analyzed how the threshold of the ganglion cells changes when nearby amacrine cells are depolarized or hyperpolarized. For each ganglion cell and amacrine cell condition, the relation between firing rate and filtered input was recorded (c.f. Method of intracellular recording). Fitting the two response curves with sigmoid functions yielded thresholds of a ganglion cell during the hyperpolarizing (*µ*_h_) and the depolarizing (*µ*_d_) current injection to the amacrine cell. The difference in thresholds (*µ*_d_ − *µ*_h_) reflects the impact of amacrine cell inputs on the response properties of the ganglion cell. We analyzed these differences as a function of the receptive field distance between the ganglion and amacrine cells. Overall, the analysis was based on current injection to 40 different amacrine cells and recordings from 144 Off ganglion cells. We note that an amacrine cell usually connects to multiple ganglion cells, and some of the ganglion cells receive inputs from multiple amacrine cells. The red and blue points shown in Fig. 6 are obtained by binning (according to RFs distance) results from 169 amacrine-to-adapting cell pairs and 32 amacrine-to-sensitizing pairs, respectively. The standard error in RFs distance (*x*-axis error) is too small to be visible in the plot.

## Acknowledgments

This research was supported by NSF CAREER Award IIS-1254123, NSF grants IOS-1556388 and IIS-1724421 (T.O.S.); 2016 Salk Women & Science Special Award, 2018-2019 Bert and Ethel Aginsky Research Scholar Award (W.M.H.); and NEI grants (S.A.B.). We thank to CNL-T members for many helpful discussions and advice.

## Supplementary

**Table S1:** Estimated noise components in the primary *ν* and modulatory *σµ* pathways for each cell pair.

| Cell pair # | Adapting |  | Sensitizing |  |
| --- | --- | --- | --- | --- |
| | $\nu_1(\sigma)$ | $\sigma_{\mu,1}(\sigma)$ | $\nu_2(\sigma)$ | $\sigma_{\mu,2}(\sigma)$ |
| A | $0.819\sigma$ | $0.281\sigma$ | $0.784\sigma$ | 0.0 |
| B | $0.671\sigma$ | $0.546\sigma$ | $0.749\sigma$ | 0.049 |
| C | $0.713\sigma$ | $0.448\sigma$ | $0.801\sigma$ | 0.025 |
| D | $0.413\sigma$ | $0.618\sigma$ | $0.545\sigma$ | 0.051 |
| E | $0.670\sigma$ | $0.598\sigma$ | $0.758\sigma$ | 0.010 |
| F | $0.604\sigma$ | $0.528\sigma$ | $0.622\sigma$ | 0.033 |
| G | $0.482\sigma$ | $0.751\sigma$ | $0.594\sigma$ | 0.013 |
| Combined fit | $0.597\sigma$ | $0.563\sigma$ | $0.685\sigma$ | 0.033 |

The dependence of noise components upon the contrast are fitted by cell pair, across the nine contrasts ( $\sigma = 12\%$ to $36\%$ in $3\%$ intervals). All the fitting parameters are in the unit of critical noise value, $\nu_c$ .

**Figure S1:**
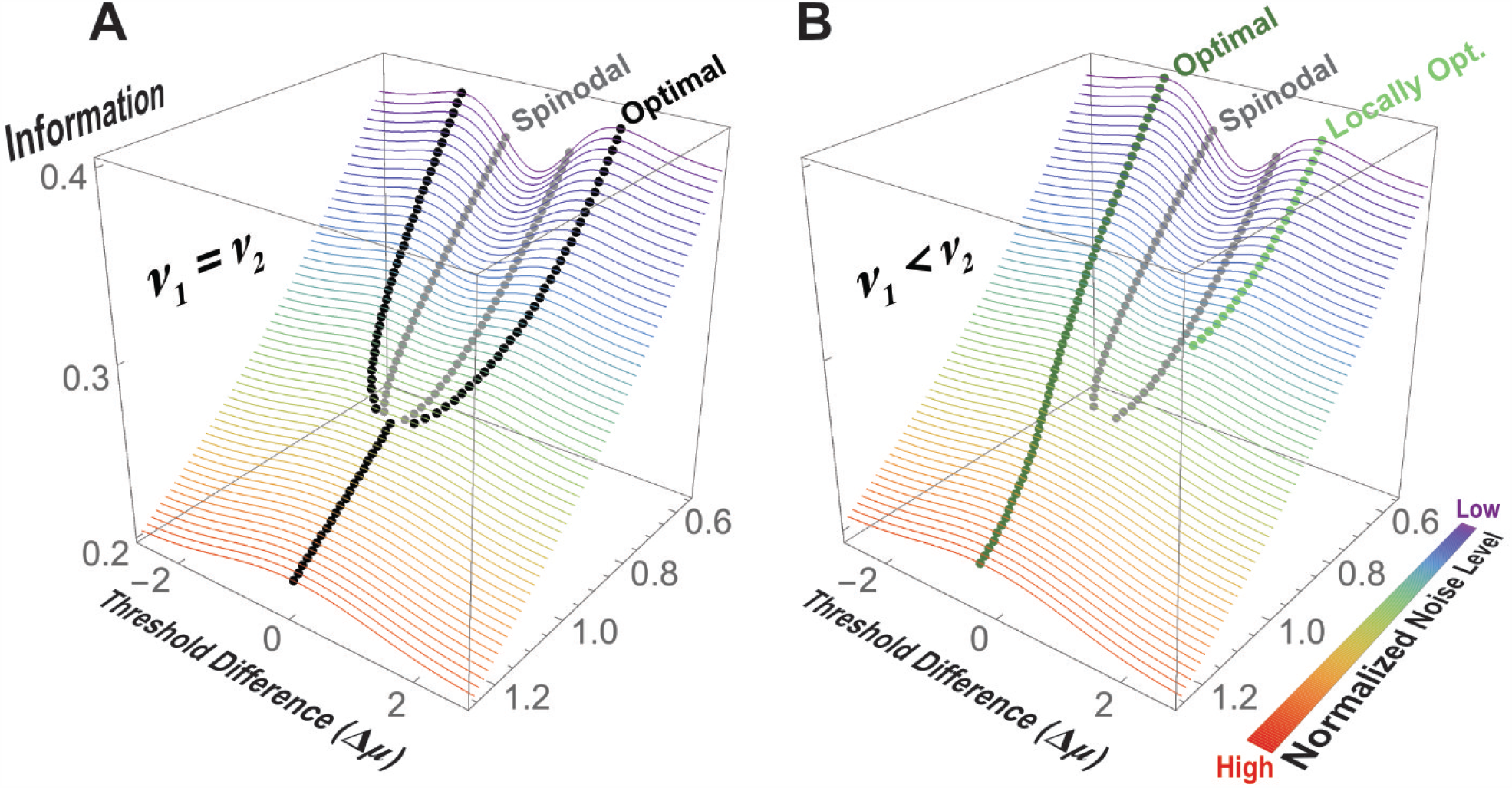
Information predicts the optimal thresholds for a pair of neurons given the values of noise level. (**A**) and (**B**) shows the information transmitted by a pair of neurons at different values of noise and thresholds. The noise level is the same for the two neurons in (**A**) and different in (**B**), ∆*ν/ν*_*c*_ = *−* 0.02, between neurons. Black and dark-green dots mark global and local information maxima, respectively. Local maxima appear when noise levels differ across neurons (**B**); otherwise the maxima are equivalent as in (**A**). Gray dots mark the inflection points, the so-called spinodal lines that delineate the regions where local maxima can be found.

**Figure S2:**
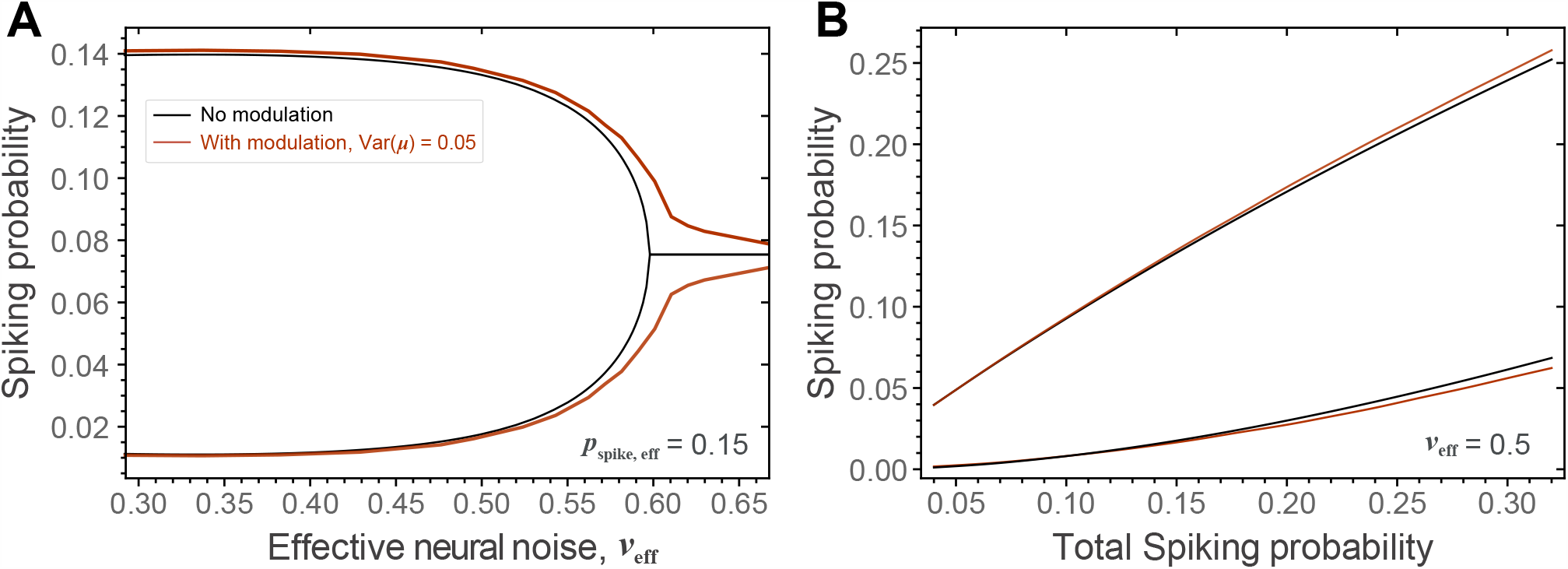
The maximally informative spike rates among neurons were similar for models with and without threshold modulation. The results shown here correspond to the analyses of the impact of threshold modulation shown in Fig. 2.

**Figure S3:**
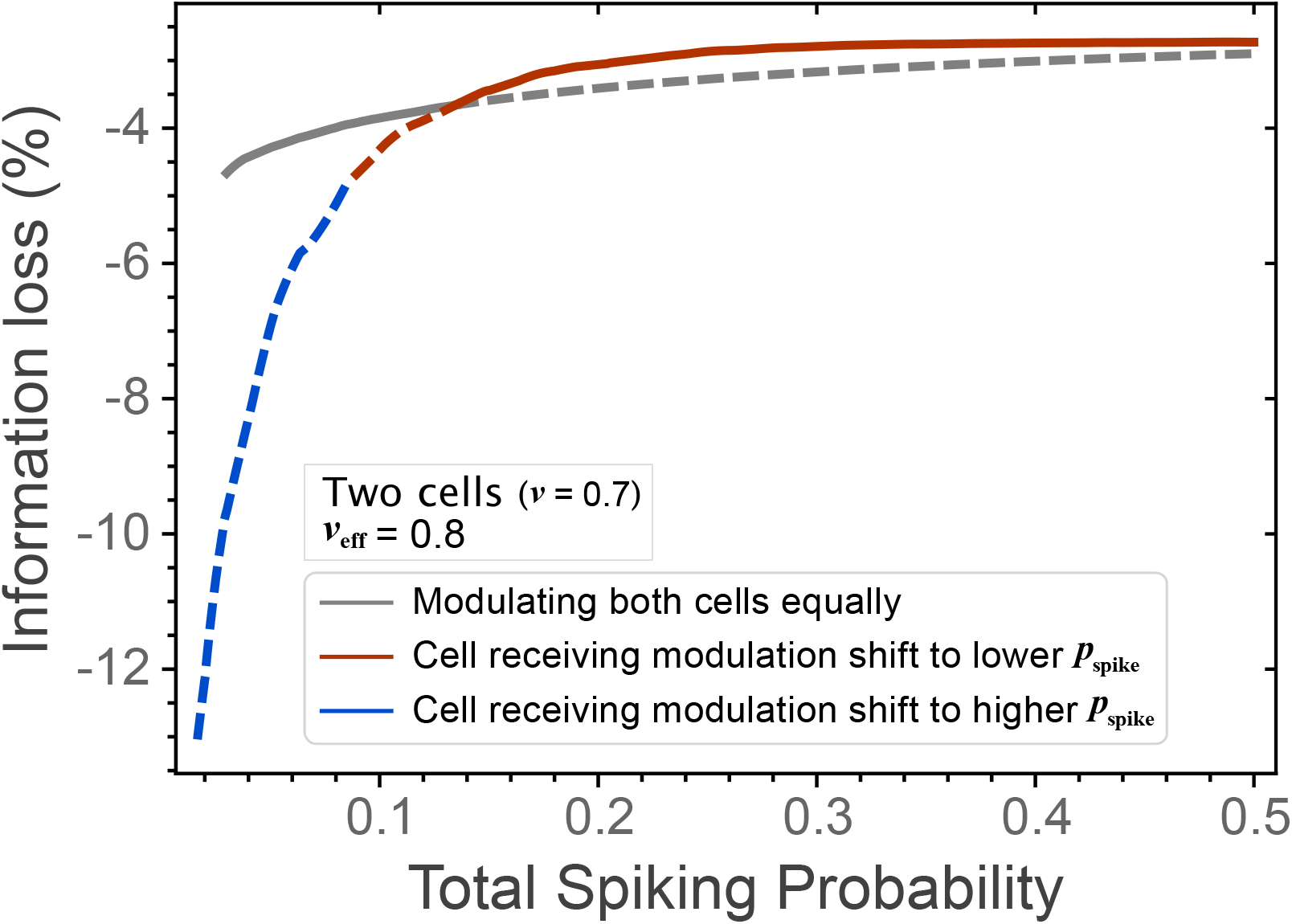
Optimal ways to apply modulation for neurons with identical spike rates depend on their rates. For neurons with low spike rates, the negative impact of threshold modulation is minimized when modulation is applied equally to both neurons (gray line). For neuron with large response rates, selective application of modulation to one of the neurons is preferred. The neuron receiving modulation shifts its threshold to decrease its response rate (red line). Curves are shown as dashed in the regimes when they become sub-optimal in terms of information transmission in the presence of threshold modulation.

**Figure S4:**
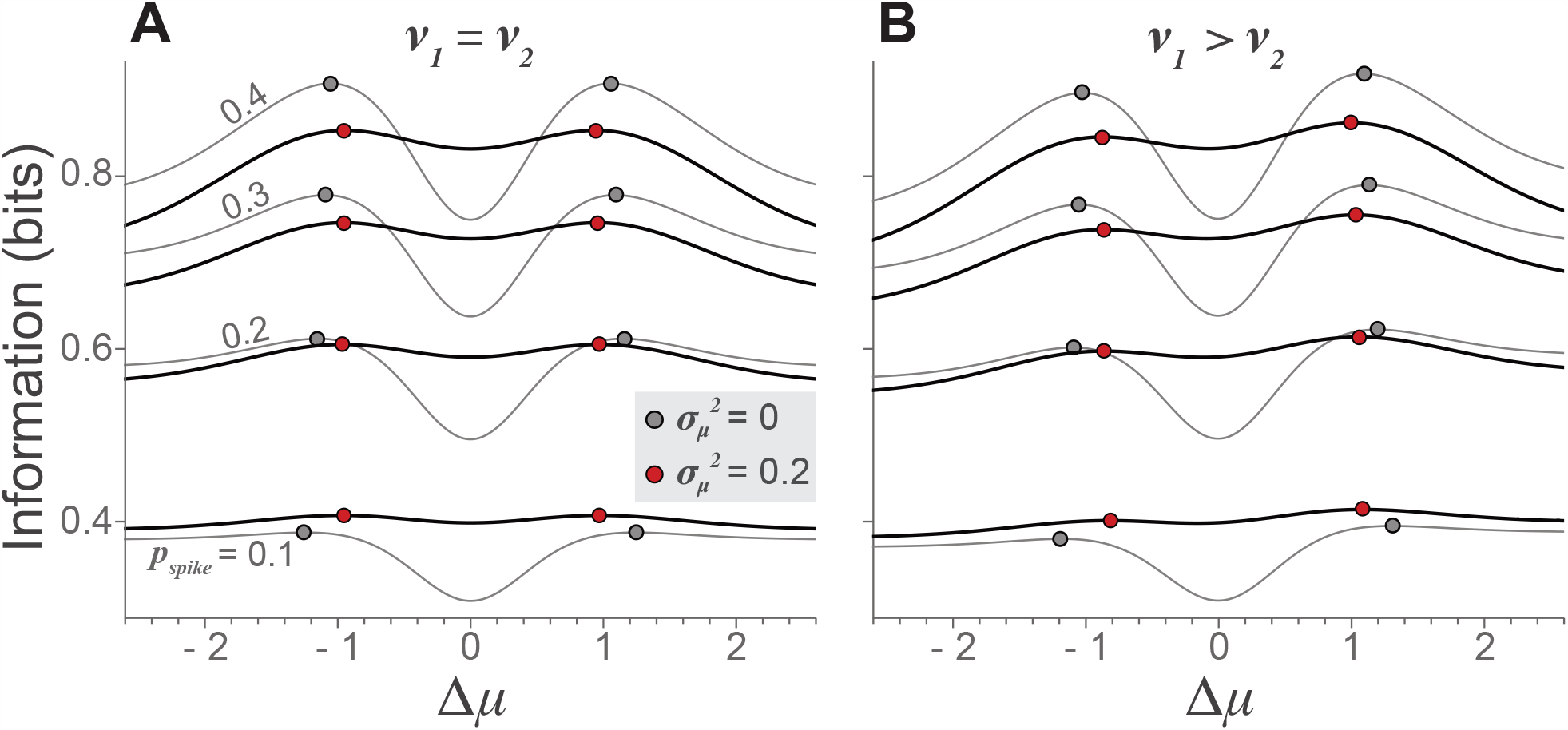
Long-term information surface. The threshold modulation effectively smooths the information surface (thin lines, gray dots mark maxima) to give rise to the long-term information surface (thick lines, red dots mark maxima).

